# A Novel Whole Tissue Explant Model of Hidradenitis Suppurativa

**DOI:** 10.1101/2024.07.19.603617

**Authors:** PE Leboit, DU Patel, JN Cohen, MI Moss, HB Naik, AE Yates, HW Harris, DM Klufas, EA Kim, IM Neuhaus, SL Hansen, RL Kyle, M Kelly, MD Rosenblum, MM Lowe

## Abstract

Hidradenitis Suppurativa (HS) is a relatively common and highly morbid inflammatory skin disease. Due to our relatively limited understanding of HS’s pathogenesis, there are currently insufficient treatment options available, and many patients’ medical needs are not being met. This is partly due to a scarcity of ex vivo human assays and animal models that accurately recapitulate the disease. To address this deficit, we have developed a whole-tissue explant model of HS to examine its pathogenic mechanisms and the efficacy of potential treatments within intact human tissue. We measured cytokine protein and RNA within whole tissue maintained in an agar-media solution, finding that IL-6 and IL-8 concentrations trended upwards in both HS explants and healthy controls, while IL-17A, IL-1β, and TNF-α exhibited increases in HS tissue alone. We also show that the explants were responsive to treatment with both dexamethasone and IL-2. Not only do our results show that this model effectively delivers treatments throughout the explants, but they also elucidate which cytokines are related to the explant process regardless of tissue state and which are related to HS tissue specifically, laying the groundwork for future implementations of this model.

## INTRODUCTION

Hidradenitis Suppurativa (HS) is an inflammatory skin disease that affects approximately one percent of the US population and disproportionately impacts women and marginalized groups (Garg et al., 2017; Straalen et al., 2022). HS is caused by a combination of genetic, endocrine, immune, and environmental factors, with specific risk factors including obesity, smoking, and a family history of the disease (Jiang et al., 2021; Seyed et a1., 2020). Patients with HS are burdened by a high frequency of comorbidities, with higher rates of inflammatory bowel diseases and mental health disorders than the general population, as well as a greater five-year mortality rate than patients with other inflammatory skin diseases (Naik and Lowes, 2019).

HS lesions typically occur around the breast, axillae, buttocks, and groin regions (Straalen et al., 2022). These lesions begin as defects in the hair follicle that then progress into follicular plugs and, eventually, to the formation of cysts. The cysts can rupture, bathing the surrounding tissue with inflammatory mediators. In some patients, these lesions become fibrotic and advance into a chronic condition (Seyed Jafari et al., 2020).

Several treatments targeting different cytokines implicated in HS immunopathogenesis have been approved or are currently undergoing clinical trials. Adalimumab (Humira), a TNF-α inhibitor, is one of two FDA-approved treatments for HS, and other systemic anti-TNF-α therapies, including infliximab, certolizumab, and golimumab, have shown promising results (Fragoso et al., 2023; Grant et al., 2010). Ruxolitinib, a topically applied JAK 1/2 inhibitor, has been found to halve concentrations of TNF-α, IL-6, IL-8, IL-1β, and CXCL3 *in vitro* in HS lesional keratinocytes and is currently being tested in two phase II clinical trials (Fragoso et al., 2023; Schell et al., 2023). Targeting the IL-17 family of cytokines and their receptors, as well as IL-23, has also reduced many patients’ symptoms in early-stage clinical trials, leading to the approval of one IL-17A antagonist, secukinumab (Fragoso et al., 2023; COSENTYX®). However, many of these treatments have variable efficacy for individual patients and have caused serious side effects in trial participants, including worsening of HS symptoms and development of irritable bowel disease, indicating a continued need for new therapeutic strategies (Fragoso et al., 2023).

HS lesions are characterized by expanded T helper 17, neutrophil, plasmacytoid dendritic cells, and plasma cell populations, as well as elevated levels of IL-1β, IL-17, IL-23, and TNF-α (Moran et al., 2023; Seyed Jafari et al., 2020). With that said, the dominant inflammatory mediators driving HS pathology have yet to be fully resolved (Zouboulis et al., 2020). This is partially due to the lack of murine and human *ex vivo* models for HS (Frew and Piguet., 2020). In recent years, several groups have been working to develop accurate, accessible *ex vivo* models for HS. Our group observed that treating *ex vivo* cultures of cells from HS lesions with TNF-α therapeutic antibodies reduces T cell activation and proliferation, mimicking results seen in patients (Lowe *et al*., 2020). Both Vossen *et al*., (2019) and Moran *et al.,* (2023) employed an explant culture model to examine the cytokine production of punch biopsies taken from HS lesions in an array of conditions. Vossen *et al*. quantified cytokine RNA within explants and cytokine protein released in the media after culture with several anti-inflammatory agents currently being used to treat HS, finding a large variation in efficacy and illuminating the need for deeper investigation into the underlying biology of patient response to different treatments. Moran *et al*. quantified cytokines released into the explant media, finding overall elevated levels of IL-1β, IL-18, IL-17A, IFNγ, and IL-36γ compared to healthy controls, with a rich heterogeneity of cytokine production between HS lesions. However, neither study measured phenotypic changes directly within tissue.

In the current study, we examine how HS pathology is affected by the explant process relative to healthy control tissue as well as how tissue responses can be altered through anti-inflammatory agents. We measure cytokine production within tissue on the RNA and protein levels. Unlike previous studies, we quantify cytokine concentrations directly within the tissue and use media infused with agar in order to maintain tissue-air interfaces and standardize the amount of tissue exposed to media. We find that the production of some cytokines, including IL-8 and IL-6, was linked to the explant state independently of disease status, while that of other cytokines, such as IL-1β, TNF-α, and IL-17A, was higher in HS biopsies than in healthy controls, both at baseline and after explantation. This novel model may allow researchers to profile the effects of different treatments on whole tissue in a more standardized manner.

## MATERIALS AND METHODS

### Ethics Statement

Discarded de-identified tissue was certified as Not Human Subjects Research per institutional guidelines. Patients providing tissue with associated clinical metadata provided written, informed consent under protocol 13-11307. The UCSF Institutional Review Board approved the proposed studies.

### Skin

24 HS lesional samples and 23 healthy controls were acquired. HS lesional skin was obtained as fresh surgical discards from patients who had undergone surgical resection or deroofing procedures. Healthy skin was obtained as fresh excess skin from reduction mammoplasty, herniorrhaphy, or Mohs surgery. All samples were dermatomed at a depth of 800 micrometers before punch biopsies were taken. Due to assay limitations, the majority of samples were exclusively used for RNA or for protein readouts.

### Explant assay

6 mm or 8 mm punch biopsies were taken from HS lesional skin. 4 mm, 6 mm, or 8 mm punch biopsies were taken from healthy control skin. Biopsies were flash frozen for protein assessments and stored at −80**°**C for future processing or placed in RNA*later* (Invitrogen, Thermo Fisher Scientific, AM7020) for RNA assessments at 4**°**C for 24 hours before being stored at −80**°**C for future processing. To prepare the explant assay medium, 3% UltraPure^TM^ Low Melting Point Agarose (Thermo Fisher Scientific, 16520050) was melted in heated PBS.

After cooling to 37**°**C in a water bath, the agar was mixed 1:1 with Immunocult: XF Cell Expansion Media (Immunocult, Stemcell, 5543916). 2 mL of explant assay medium was plated in individual wells of a 24-well plate and allowed to solidify. Baseline samples were preserved immediately, and explant samples were placed on top of the supportive agar-based medium and cultured for 24 hours at 37**°**C in a 5% CO2 incubator.

### *Ex vivo* assay method

To produce single cell suspensions, explants were enzymatically digested directly following incubation using collagenase IV (1000 U/mL, Worthington, LS004186) and DNAse (200 μg/mL) for two hours in RPMI + 10% FBS. Samples were then plated in a 96-well U-bottom plate, with 300,000 cells allocated to each well, and cultured for two to three days at 37**°**C in a 5% CO2 incubator. Directly after incubation, supernatants were collected and banked at −20**°**C for protein quantification. Cytokine concentrations were then quantified via Human Cytokine Panel A 48-Plex Discovery Assay® Tissue-Cell Culture (Luminex, Eve Technologies, Calgary, AB, Canada).

### Compounds

Dexamethasone (Sigma-Aldrich, D4902-25MG) was prepared at a 10000x concentration in methanol and dosed at 10 μM. Cpd A (Sitryx Therapeutics, Oxford, UK), a fluorescent metabolic inhibitor with modest hydrophilic properties (logD =0.9 at pH 7.4, calculated logP = 2.7), was prepared at 1000x concentrations in DMSO and dosed at 31.62 μM. Proleukin IL-2 (UCSF Pharmacy) was obtained at 500,000 IU/mL and aliquoted for storage at - 80**°**C before use; it was dosed at 3000 IU/mL. All compounds were resuspended within the explant assay medium prior to its solidification.

### Protein quantification

Biopsies were homogenized in Eve Technologies’ recommended protease inhibitor buffer (20 mM Tris HCl, 0.5% Tween 20, 150 mM NaCl, cOmplete™, Mini, EDTA-free Protease Inhibitor Cocktail (Sigma-Aldrich, 11836170001) in diH_2_0) using the Precellys® Evolution Touch Homogenizer for three 20-second spins with two 30-second pauses in between (6800 rpm), and supernatants were collected following centrifugation for 15 minutes (2000 xg). Cytokine concentrations of GM-CSF, IFN-γ, IL-1β, IL-2, IL-4, IL-5, IL-6, IL-8, IL-10, IL-12p70, IL-13, IL-17A, IL-23, and TNF-α were then quantified via Human High Sensitivity 14-Plex Discovery Assay® Tissue Homogenate (Luminex, Eve Technologies, Calgary, AB, Canada). For the dexamethasone experiments, cytokine concentrations were quantified via the Human Cytokine Panel A 48-Plex Discovery Assay® Tissue Homogenate (Luminex, Eve Technologies, Calgary, AB, Canada). Cytokine concentrations were normalized to biopsy weight.

### Messenger RNA quantification

RNA was extracted using the RNAeasy Fibrous Tissue Mini Kit (Qiagen, 74704), and cDNA libraries were assembled using the High-Capacity cDNA Reverse Transcription Kit (ThermoFisher, 4368814) and the Veriti 96 Well Thermal Cycler (Applied Biosystems, Waltham, MA). Concentrations of cDNA were 100 ng/uL when possible. cDNA concentrations were standardized within individual donor experiments. Using the specified primers, real-time quantitative polymerase chain reaction (qPCR) was performed to quantify transcripts of *IL1B* (Thermo Fisher, Hs01555410_m1), *IL17A* (Thermo Fisher, Hs00174383_m1), *TNF* (Thermo Fisher, Hs00174128_m1), *IL6* (Thermo Fisher, Hs00985639_m1), *CXCL8* (Thermo Fisher, Hs99999034_m1), *CSF2* (Thermo Fisher, Hs00929873_m1), *IL23* (Thermo Fisher, Hs00372324_m1), *CD3E* (Thermo Fisher, Hs01062241_m1) and *PTPRC* (Thermo Fisher, Hs04189704_m1). Primers were added to Taqman^TM^ Fast Advanced Master Mix for qPCR (Thermo Fisher, 4444554) and then combined with cDNA in a MicroAmp^TM^ Fast Optical 96-well Reaction Plate, 0.1 mL (Applied Biosystems, 4346907) prior to being run on the StepOnePlus Real-Time PCR System (Applied Biosystems, Waltham, MA). All measurements were normalized using the housekeeping genes *GUSB* (Thermo Fisher, Hs99999908_m1), *GAPDH* (Thermo Fisher, Hs99999905_m1), and *HPRT* (Thermo Fisher, Hs99999909_m1).

### Frozen sectioning

Explants were embedded in OCT and frozen using liquid nitrogen. They were then sectioned at a thickness of 20 µm and imaged using the DAPI 10xPLFL-DRY channel to measure treatment fluorescence and the Spectrum Green 10xPLFL-DRY channel for background normalization on the Aperio Versa 8 (Leica Biosystems, South San Francisco, CA).

### Histology

Explants were preserved for at least 24 hours in 10% formalin (Fisher Chemical, SF100-20) before paraffin embedding. The UCSF Dermatopathology Service performed embedding, sectioning (3 µm), and staining with hematoxylin and eosin. Brightfield imaging was performed with the Aperio Versa 8 (Leica Biosystems, South San Francisco, CA).

### Flow cytometry

To produce single-cell suspensions, explants were enzymatically digested directly following incubation using collagenase IV (1000 U/mL, Worthington, LS004186) and DNAse (200 μg/mL) for two hours in RPMI + 10% FBS. Samples were then stained extracellularly using Ghost Dye^TM^ Violet 510 (Tonbo Bioscience, 13-0870-T100), Brilliant Violet 711^TM^ anti-human CD45 Antibody (BioLegend, 304050), PE-CF594 Mouse anti-Human CD4 (BD Horizon, 562281), CD3 Monoclonal Antibody APC-eFluor^TM^ 780 (Invitrogen, 47-0036-42), Brilliant Violet 650^TM^ anti-human CD8a Antibody (BioLegend, 301042), and BD^TM^ APC Mouse Anti-Human CD25 (BD Biosciences, 340939). Fixation and permeabilization was performed using the eBioscience^TM^ Intracellular Fixation & Permeabilization Buffer Set (Invitrogen, 88-8824-00). Intracellular staining was performed using the FOXP3 Monoclonal Antibody, eFluor^TM^ 450 (Invitrogen, 48-4776-42). Samples were then run on an LSRFortessa™ X-20 Cell Analyzer (BD Biosciences, Milpitas, CA) and analyzed using FlowJo™ v10.10 Software (BD Life Sciences, Franklin Lakes, NJ).

### Statistics

To analyze transcriptomic differences between healthy and HS biopsies at baseline and following culture in media, mixed effects analysis and Sidak’s multiple comparison test were performed for each cytokine. To analyze protein level differences between healthy and HS biopsies at baseline and following culture in media, an ordinary one-way ANOVA and Sidak’s multiple comparisons test were performed for each cytokine. To analyze differences between healthy and HS *ex vivo* cultures, Welch’s t-test was performed for each cytokine. To analyze the effects of dexamethasone on cytokine transcript levels, an ordinary one-way ANOVA with repeated measures and Sidak’s multiple comparisons test were performed for each cytokine. To analyze the effects of IL-2, Welch’s t-test was performed comparing flow cytometry results from samples cultured in media with added IL-2 to those cultured in control media. All the above analyses were conducted using GraphPad Prism version 10.2.3 for macOS (GraphPad Software, Boston, Massachusetts USA, www.graphpad.com). To analyze the effects of dexamethasone on cytokine protein concentrations, a linear mixed-effects analysis was performed for each cytokine in RStudio (v4.0.5; RStudio Team, 2021) using the lme4 package (v1.1.35.1; Bates et al., 2023). Condition and surgical procedure were input as the fixed effects, while donor and replicate, as well as by-donor and by-replicate random slopes for the effect of treatment, were input for the random effects.

## RESULTS

We established an explant model of intact skin tissue with consistent tissue volume and reagent exposure by first subjecting HS skin to a consistent depth *via* dermatome and then creating equivalently-sized punch biopsies (Figure 1A). After 24 hours of culture, microscopic evaluation showed that explanted HS skin had decreased perivascular and interstitial lymphocytic infiltrates and a zone of keratinocyte necrosis in the upper epidermis as well as in the eccrine secretory glands compared to baseline (Figure 1B). Similarly, explanted healthy skin cultured in media also showed a degree of keratinocyte necrosis in these micro-anatomic compartments, which most likely reflects ischemic changes. Enzymatic digestion of explants post-culture produced an average of 1.43 million cells (viability of 65%) (Figure 1C), and quantitative PCR analysis of the common leukocyte receptor, *PTPRC* (CD45), and the epsilon chain of the T cell receptor-CD3 complex, *CD3E*, showed good retention of immune populations (Figure 1D). Next, we determined the kinetics of penetration of a small molecule through the full thickness of the explant. To this end, we used Cpd A, an auto-fluorescent small molecule with modestly hydrophilic properties (logD =0.9 at pH 7.4, calculated logP = 2.7). Cpd A was dissolved in the explant media, and explants were cultured for 4, 24, or 48 hours. Imaging revealed drug exposure began by 4 hours and saturated tissue by 24 hours (Figure 1E).

**Figure 1.**
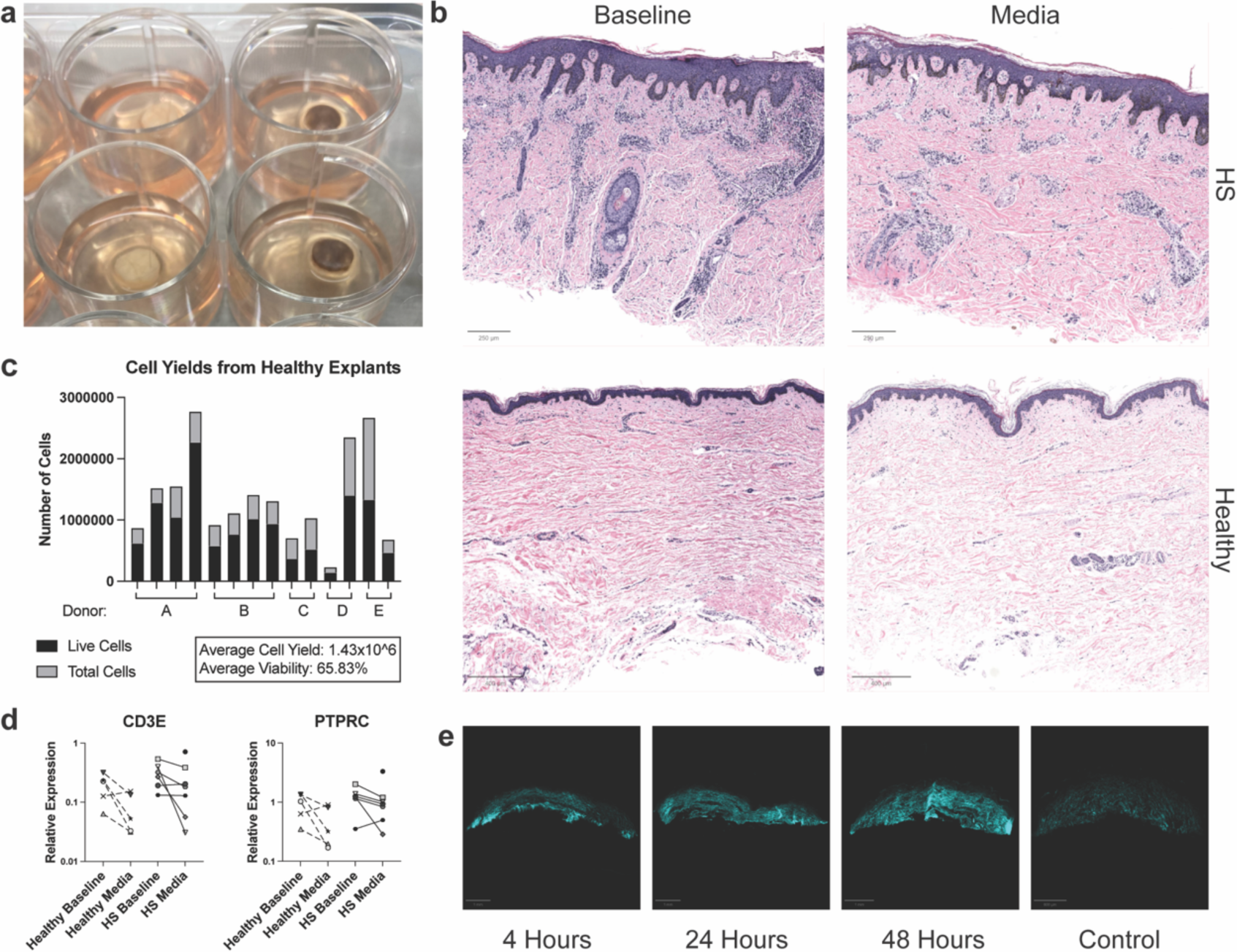
Explants cultured for 24 hours maintain live immune cell populations and exhibit complete drug penetration. **(A)** HS explants cultured for 24 hours. **(B)** H&E-stained sections of HS (250 µm scale bar) and healthy control (400 µm scale bar) explants (representative of 3 independent healthy samples and 2 independent HS samples). **(C)** Cell yields following 24-hour culture (Each bar represents one explant). **(D)** Quantitative reverse-transcriptase polymerase chain reaction (RT-qPCR) analysis of CD3E and PTPRC (CD45) in healthy (n=5) and HS explants (n=7) at baseline and after culture relative to housekeeping gene expression (ns, mixed-effects analysis with Šídák’s multiple comparisons test). Each shape represents one donor (averaged for 1-4 explant tissues, solid line = HS donor, dashed line = healthy control). **(E)** Frozen sections of healthy explants cultured with Cpd A for the specified time (representative of 3 independent samples).

Therefore, it was determined that the 24-hour incubation time maximized treatment delivery of a small molecule while minimizing cell loss.

We next examined IL-8, IL-6, GM-CSF, IFNγ, IL-1β, TNF-a, IL-17A, and IL-23A, which have previously been shown to be important to the immunopathogenesis of HS lesions, at baseline and within the explant model (Kelly et al., 2015; Straalen et al., 2022). We compared HS to healthy skin tissue RNA and protein at baseline and throughout culture (Figure 2A and B). At baseline, concentrations of IL-17A protein were significantly increased in HS relative to healthy tissue. Baseline protein concentrations of TNF-α and IFNγ trended higher in HS than in healthy tissue. Explant culture resulted in upward trends in IL-8 and IL-6 on the RNA and protein levels in both healthy and HS explants, which reached significance in HS tissue RNA. Additionally, during incubation, several cytokines that have been targeted by HS therapeutics clinically—TNF-α, IL-17A, and IL-1β—increased more in HS than they did in healthy control tissue. This elevation was significant for IL-1β on the protein and RNA levels. After explant culture, the final protein concentrations of IL-1β, TNF-α, and IL-17A were all significantly higher in HS than in healthy skin.

**Figure 2.**
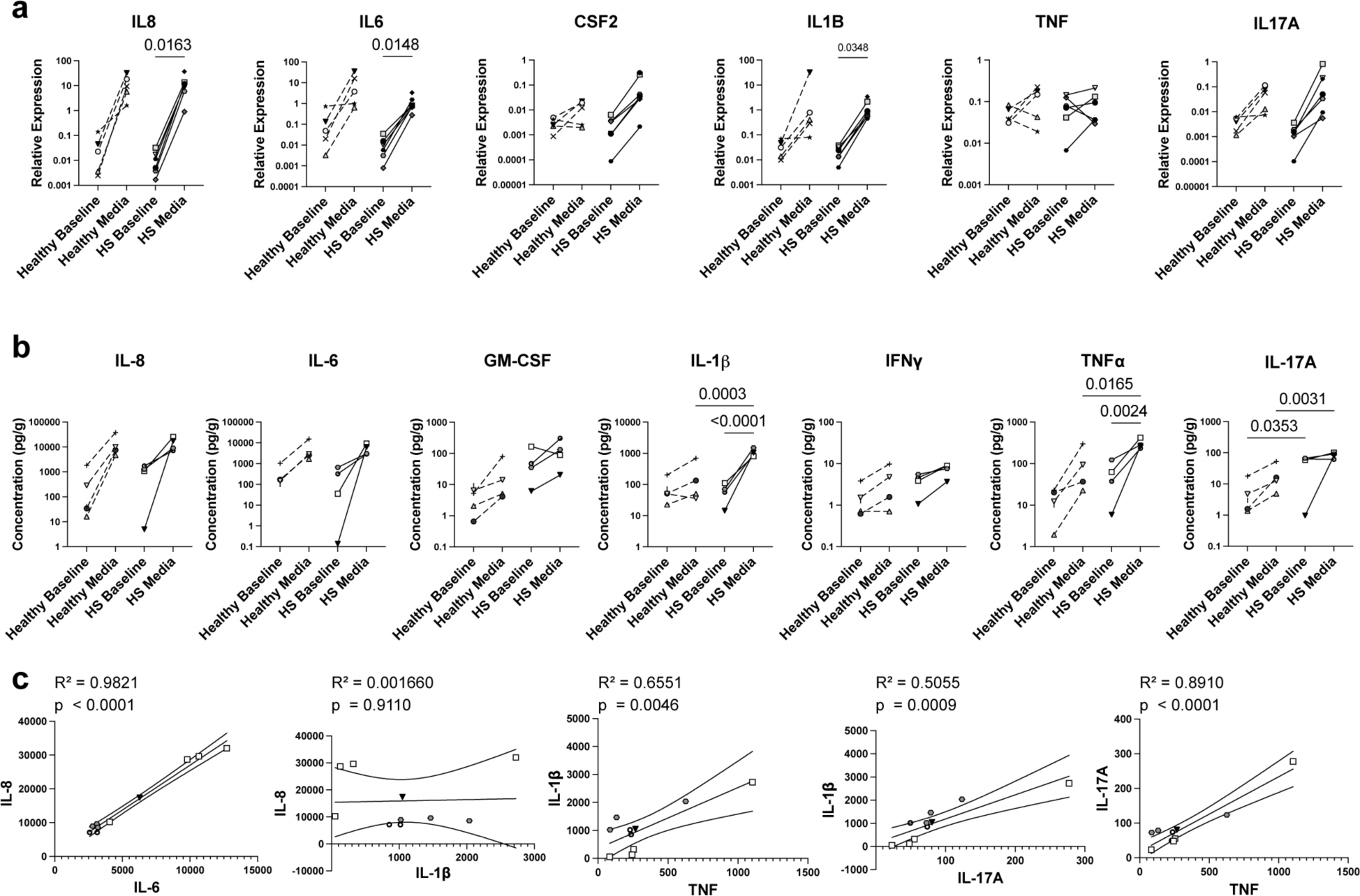
Concentrations of IL-6 and IL-8 trend higher across disease state, while upward trends in the concentrations of IL-17A, IL-1B, and TNF-a are specific to HS. **(A)** RT-qPCR analysis of cytokines relative to housekeeping gene expression in healthy (n=5) and HS (n=6) explants at baseline and following 24-hour culture (mixed-effects analysis with Šídák’s multiple comparisons test). Each shape represents one donor (averaged for 1-4 explants, solid line = HS donor, dashed line = healthy control). **(B)** Cytokine protein concentration of healthy (n=5) and HS (n=4) explant tissue quantified via Luminex Assay (ordinary one-way ANOVA with Šídák’s multiple comparisons test). Each shape represents one donor (averaged for 1-4 explants, solid line = HS donor, dashed line = healthy control). **(C)** Simple linear regression of cytokine protein concentrations (pg/g) in HS explants (n = 4) following 24-hour culture in media. Each shape represents one donor; each symbol represents one explant tissue.

In an attempt to understand the relationship between these cytokine trends, we examined the correlations between cytokine levels in HS explants cultured in media (Figure 2C). IL-8 and IL-6 protein concentrations were very strongly correlated, while IL-8 and IL-1β protein concentrations were very weakly correlated. In contrast, IL-1β was moderately correlated with IL-17A protein and strongly correlated with TNF-α protein. TNF-α and IL-17A protein concentrations were very strongly correlated. These findings are consistent with our observations that IL-6 and IL-8 responses are associated with explant culture, while IL-1β, TNF-α, and IL-17A protein production may be more related to tissue inflammation observed in HS skin. In opposition to protein, RNA correlations were generally more weakly related for these cytokines (data not shown).

Next, we analyzed cytokine concentrations from *ex vivo* single-cell suspension cultures of HS and healthy skin (Figure 3) to understand how the explant model compares with previously studied dissociated cell systems. As in the explant assay, IL-8 and IL-6 proteins were concentrated in both healthy and HS supernatants following dissociated cell culture, though IL-8 was significantly higher in HS than in healthy controls. Additionally, protein concentrations of TNF-α and IL-17A were significantly increased in the HS *ex vivo* culture assay. The difference in concentrations of IL-1β between HS and healthy single-cell cultures trended towards significance, while IFNγ levels were fairly similar between conditions. Unlike the explant assay, GM-CSF was significantly higher in HS than in healthy controls following single cell culture. Taken together, these data suggest that HS and healthy tissue behavior in the whole-tissue explant model demonstrates several consistencies with other assay modalities.

**Figure 3.**
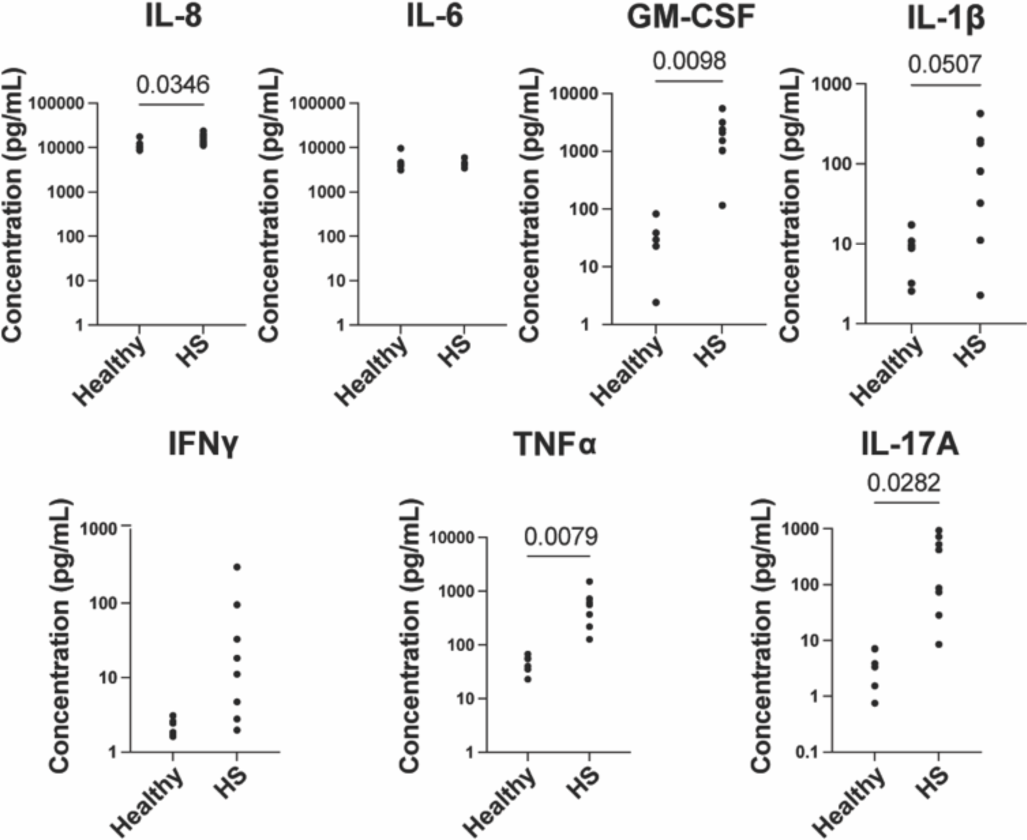
Explant cytokine concentrations are consistent with those from single-cell suspension cultures. Cytokine protein concentrations from the supernatants of healthy (n = 7) and HS (n = 8) single-cell *ex vivo* cultures after 48 hours quantified via Luminex assay (Welch’s T test). Each dot represents one healthy or HS donor (averaged for 3 single-cell suspension wells).

Finally, we tested whether therapeutic compounds are capable of altering the cytokine production or immune cell activity of these explants within 24 hours. HS explants treated with dexamethasone showed a reduction in several cytokines on the RNA level (Figure 4A). *IL6* decreased significantly when treated with 1 µM and 10 µM dexamethasone. *CXCL8* followed a similar trend for both conditions. Meanwhile, the abundance of *IL1B*, *CSF2* (GM-CSF), and *IL23* transcripts all greatly decreased when treated with 10 µM dexamethasone. Since treatment with dexamethasone appeared to affect cytokine production, an expanded Luminex panel was used to examine changes in cytokine protein concentration in response to treatment. Protein concentrations of IL-1β, CXCL1, CCL3, and PDGF-AA all significantly decreased when treated with 10 µM dexamethasone, while concentrations of IL-8 and IL-17A both trended downwards (Figure 4B). In healthy tissue, treatment with IL-2, which directly induces expression of its own receptor (Sereti et al., 2000), significantly increased the percentage of total CD4 and effector T cells expressing CD25 (the high-affinity IL-2Rα subunit) (Figure 4C). Additionally, the frequency of Tregs, which are highly dependent on IL-2 in lymphoid organs, increased significantly as a proportion of CD4 T cells (Figure 4C). Taken together, these data show that cytokine expression and CD25 protein production can be effectively altered using our explant model.

**Figure 4.**
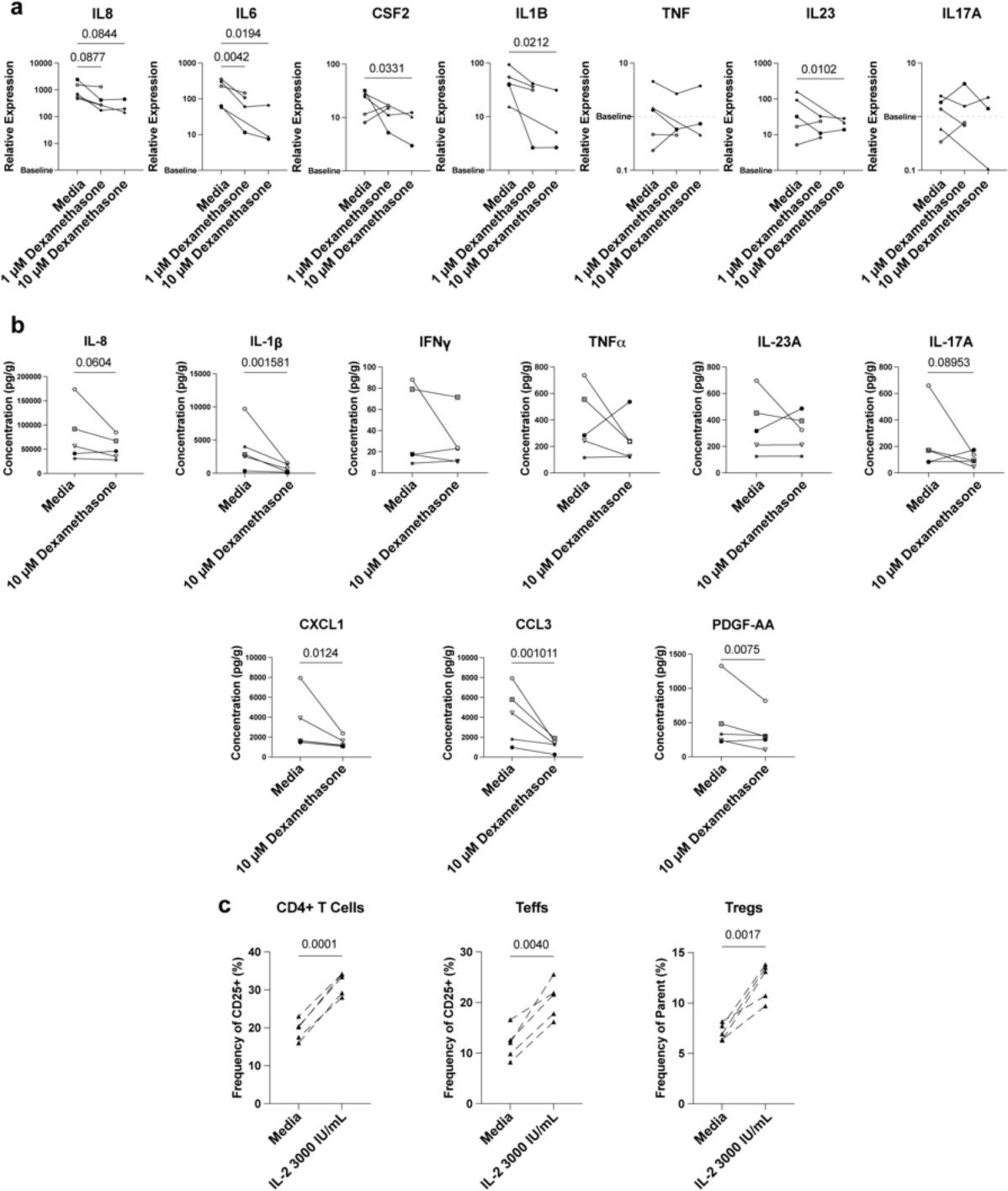
Explants cultured in media with added treatment for 24 hours showed significant differences on both the RNA and protein levels. **(A)** RT-qPCR analysis of cytokines relative to baseline in HS explants (n=5) after 24-hour culture in the specified condition (mixed-effects analysis with Šídák’s multiple comparisons test). Each shape represents one donor (averaged for 1-4 explant tissues). **(B)** Cytokine protein concentration of HS explant tissue (n=5) at baseline or after 24-hour culture in the specified condition quantified via Luminex Assay (linear mixed-effects analysis). Each shape represents one donor (averaged for 1-3 explant tissues). **(C)** Healthy explants (n=5) were cultured in media with or without 3000 IU/mL IL-2 for 24 hours and analyzed by flow cytometry (Welch’s t test). Each symbol represents one donor (averaged for 1-2 explant tissues).

## DISCUSSION

New ‘industry grade’ models are greatly needed for mechanistically probing HS pathology and testing novel therapeutic interventions. The shortage of models that accurately predict patients’ responses to treatment has complicated drug development. Therefore, more models, like this one, that use intact patient tissue and bridge the gap between *in vitro* responses and those occurring in patients are greatly needed. Given the substantial interpatient variability inherent to primary tissue samples of HS, assay standardization has been critical for experimental performance. Several steps have been taken to standardize our model: 1) utilization of dermatomed skin in all samples to achieve uniform thickness; 2) using consistent punch biopsies sizes within donors; and 3) ensuring consistent treatment exposure by infusing the media with agar. Cytokine concentrations from the whole-tissue explant model were found to be relatively consistent with our previous single-cell suspension assay in which we observed responses to clinically approved therapeutic interventions. The explant model offers the preservation of inflammatory structures during therapeutic interventions as well as an increased ability to assess cell-cell interactions and changes *via* microscopy.

During the development of this model, it was observed that some cytokines examined— IL-6 and IL-8—were elevated after culture regardless of disease state, while other inflammatory mediators—IL-1β, TNF-α, and IL-17A—increased more in HS than in healthy explants. IL-1β, TNF-α, and IL-17A may be specifically increased due to cells and pathways selectively activated in HS tissue. Moran *et al*. found that HS explants secreted significantly more IL-1β, IL-17A, and IFNγ into the media relative to healthy controls. Meanwhile, consistent with past explant studies, transcripts and protein concentrations of IFNγ and IL-1β did not increase in healthy tissue (Niel et al., 2020). Interestingly, in our model, IFNγ protein concentration was not found to be higher in HS explants than in healthy explants. As there is likely a mediator affecting the types of cells and cytokines that migrate into the culture media, this discrepancy could be due to the difference in where the cytokines were collected from (media versus within the tissue) or due to variability in cytokine production between patient cohorts. HS lesions have been shown to have higher site-to-site and patient-to-patient variability than other inflammatory skin diseases (Straalen et al., 2022).

IL-6 and IL-8 consistently increased during culture regardless of disease state, implying that normal physiological responses are also at play within the explant assay. Previous literature has shown that IL-6 and IL-8 play major roles in wound healing responses, indicating explant culture may invoke tissue damage responses in all patient samples (Johnson et al., 2020; Rennekampff et al., 2000). Moreover, IL-17A, which has also been shown to play a role in wound healing, while significantly higher in HS tissue, also trended upward in healthy explant culture (Nirenjen et al., 2023). There were high correlations between certain cytokines within our explant system, indicating that cellular mediators may be co-regulated, share cellular sources, or feedback on each other. Previous work has shown that IL-6 can decrease the concentration of TNF-α, which may be one factor contributing to the relatively low concentrations of TNF-α observed in this study in both HS and healthy skin (Paquet and Piérard, 2009). Additionally, in prior reports of explant experiments, TNF-α responses were relatively more delayed within healthy tissue, allowing us to capture acute increases only within HS lesional tissue (Ashcroft et al., 2012; Niel et al., 2020).

Dexamethasone treatment resulted in striking effects on cytokines at both transcriptional and protein levels, while IL-2 affected CD25 protein expression. Dexamethasone resulted in a reduction in *IL6*, *IL1B*, *CSF2*, and *IL23* transcripts, as well as a reduction in IL-1β, CXCL1, CCL3, and PDGF-AA concentrations. It also resulted in a strong downward trend for *CXCL8* transcripts and IL-8 and IL-17A protein concentrations. IL-1β has been previously shown to have a strong response to dexamethasone, as has GM-CSF (Abraham et al., 2006; Noreen et al., 2021; Paquet and Piérard, 2009). Previous studies have also found dexamethasone to inhibit the production of TNF-α and IFNγ, but decreases were not observed in our model (Abraham et al., 2006; Noreen et al., 2021). However, TNF-α transcript was relatively lower than other cytokines within our model, which may have limited the detectable effect of dexamethasone. Additionally, since TNF-α and IFNγ protein concentrations did trend downwards in most donors, it is possible that a difference would be observed with a larger sample size. Our data examining the effects of IL-2 treatment indicates altering cell receptor expression is also feasible and that individual targets of interest must be validated as functionally malleable within this system.

There are several limitations to this explant model. All explants were dermatomed to a thickness of 800 micrometers to allow for adequate nutrient and compound delivery, meaning cytokine activity in the lower dermis was not examined. Additionally, the inflammatory state of the explant is affected to some extent by processing, as seen in changes observed in explant samples versus baseline (Figure 2A, 2B). Finally, due to limited tissue size, RNA extraction and protein isolation could not be performed on the same explant, meaning that, for the most part, donors had to be allocated to one analysis or the other. This prevented the comparison of a cytokine’s RNA concentration with its protein concentration within the same explant. When implementing this model, we have found it advantageous to have several replicates in each condition, as cytokine expression can vary between explants. Nevertheless, due to our results’ alignment with those of previous *ex vivo* single-cell cultures of HS and whole-tissue explant assays, as well as our observed treatment responses, we believe that this assay will be a useful tool for studying the general pathology of and therapeutic responses in HS.

## Acknowledgments

We acknowledge Sitryx Therapeutics for provision of tool compounds. We acknowledge the UCSF Parnassus Flow CoLab (RRID: SCR_018206), which is supported in part by the following grants: NIH P30 DK063720 and NIH S10 1S10OD01822-01.

